# Spontaneous cortical activity transiently organises into frequency specific phase-coupling networks

**DOI:** 10.1101/150607

**Authors:** Diego Vidaurre, Laurence T Hunt, Andrew J. Quinn, Benjamin A.E. Hunt, Matthew J. Brookes, Anna C. Nobre, Mark W. Woolrich

## Abstract

Frequency-specific oscillations and phase-coupling of neuronal populations have been proposed as an essential mechanism for the coordination of activity between brain areas during cognitive tasks. To provide an effective substrate for cognitive function, we reasoned that ongoing functional brain networks should also be able to reorganise and coordinate in a similar manner. To test this hypothesis, we use a novel method for identifying repeating patterns of network dynamics, and show that resting networks in magnetoencephalography are well characterised by visits to short-lived transient brain states, with spatially distinct power and phase-coupling in specific frequency bands. Brain states were identified for sensory, motor networks and higher-order cognitive networks; the latter include a posterior higher-order cognitive network in the alpha range (8-12Hz) and an anterior cognitive network in the delta/theta range (1-7Hz). Both higher-order cognitive networks exhibit especially high power and coherence, and contain brain areas corresponding to posterior and anterior subdivisions of the default mode network. Our results show that large-scale cortical phase-coupling networks operate in very specific frequency bands, possibly reflecting functional specialisation at different intrinsic timescales.

## Introduction

Efficient neuronal coordination between regions across the entire brain is necessary for cognition (Salinas and Sejnowski, 2001; Varela et al., 2001; Buschman and Miller, 2007; Siegel et al., 2012). A proposed mechanism for such coordination is oscillatory synchronisation; that is, populations of neurons transmit information by coordinating their oscillatory activity with the oscillations of the receptor population at certain frequencies. Furthermore, different frequencies, or, more generally, different oscillatory patterns, subserve different functions (Buzsáki and Draguhn, 2004). At the same time, phase-coupling, or equivalently coherence, between neuronal populations in specific frequency bands has been proposed as a mechanism for regulating the integration and flow of cognitive content (Fries, 2005; Fries, 2015; Engel et al., 2013, Marzetti et al., 2013). The role of phase-coupling at distinct frequencies has also been demonstrated in tasks at the large-scale, where task-relevant information is effectively transmitted through phase-locking between separate cortical regions (Palva et al., 2005; Hipp et al., 2011; Fries 2015; Fell and Axmacher, 2011).

Using fMRI, it has been shown that large-scale networks activated in tasks are also spontaneously recruited in the resting state (Smith et al., 2009). These networks have previously been shown to have distinct band-limited power in electroencephalography (EEG) and magnetoencephalography (MEG) (Laufs el al., 2003; Brookes et al., 2011; de Pasquale et al. 2010; Ganzetti and Mantini, 2013; Hipp and Siegel, 2015; Colclough et al. 2017). If these spontaneously occurring networks are to provide an effective substrate for cognitive processes, then they might also be expected to exhibit the same fast changing phase-coupling activity observed in tasks (Womelsdorf et al., 2007; Bosman et al., 2012; Fries, 2015). However, the evidence for frequency specific phase-coupling in spontaneous activity at timescales associated with fast cognition is limited.

Here, we propose that cortical activity at rest can be described by transient, intermittently reoccurring events in which large-scale networks activate with distinct spectral and phase-coupling features. To identify the possible presence of these events, we use a new analysis approach based on the Hidden Markov Model (HMM; Rabiner, 1989). For the first time, this allows for the identification of brain-wide networks (or brain states) characterised by specific patterns of power *and* phase-coupling connectivity, which, crucially, are spectrally-resolved (i.e. power and phase-coupling are defined as a function of frequency). These patterns are also temporally-resolved, meaning that the method provides a *probabilistic* estimation of when the different networks are active (see **Fig. 1a**). Notably, applying this approach to resting MEG recordings of healthy human subjects revealed the distinct temporal and spectral properties of anterior versus posterior regions of the default mode network. The joint description of the spectral, temporal and spatial properties of ongoing neuronal activity provides new insight into the large-scale circuit organization of the brain (Woolrich and Stephan, 2013).

## Results

Using concatenated MEG resting-state data from 55 subjects, mapped to a 42-region parcellation using beamforming (Van Veen et al., 1997; Woolrich et al., 2011) with reduction of spatial leakage in order to diminish the effects of volume conduction (Colclough et al, 2015), we identified 12 HMM states using a novel approach that we refer to as *time-delay embedded HMM*. Essentially, this technique finds, in a completely data-driven way, recurrent patterns of network (or HMM state) activity. Each HMM state has parameters describing brain activity in terms of power covariations and, crucially, phase-coupling between every pair of regions. The method provides information that is both spectrally-resolved (power and phase-coupling are defined as a function of frequency) and temporally-resolved (different networks are described as being active or inactive at different points in time). Importantly, while the spatial and spectral description of the states is common to all subjects, each subject has their own *state time course*, representing the probability of each HMM state being active at each instant (see Methods for further details). See **Fig. 1** for a graphical example, and **Fig. SI-1** for an illustration of the entire pipeline.

**Fig. 1.**
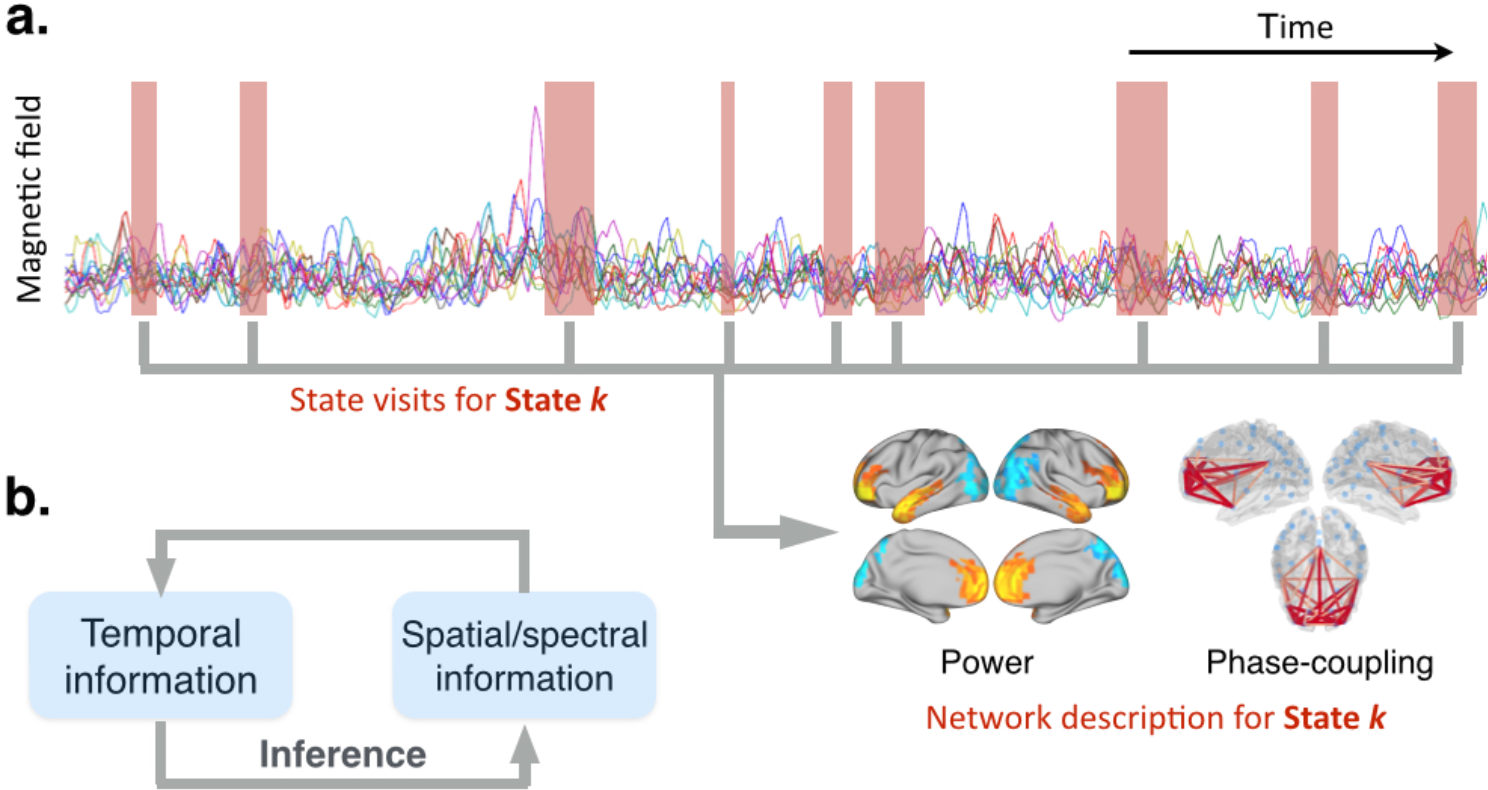
Schematic illustration of the method. **(a)** Each state is defined as having its own distinct temporal, spatial and spectral characteristics. The temporal information is given by when the state is active (red boxes). The spatial and spectral descriptions of the power maps and phase-coupling networks are contained in the parameters of each state. **(b)** Schematic of the iterative model inference. The state parameters are estimated using those segments of the data for which the state is currently estimated to be active. In turn, the estimation of when a state is active is based on the how well each state can explain each time point (i.e. according to the current estimate of the spatial/spectral state parameters).

### The states exhibit specific phase-locking connectivity

**Fig. 2** shows spatial maps of power and phase-coupling connectivity, both averaged across a wideband frequency range (1-30Hz), for four of the 12 estimated states. The maps are thresholded for ease of visualisation. **Fig. SI-2** shows the remaining eight states, four of which exhibit reduced power and connectivity relative to the grand average. The power maps and phase-coupling connectivity of each state tend to be (although not exclusively) bilateral, with strong increases in power tending to (although not exclusively) accompany increases in phase-locking. We refer to two of the states (left) as being “higher-order cognitive”, in accordance with the brain areas they incorporate and previous literature (Smith et al, 2009; Svoboda et al., 2006; van Overwalle, 2009; Mason et al., 2007). The other two states (right) correspond well to visual and motor systems. The two higher-order cognitive networks involve regions that together form the default mode network (DMN). This affords the interpretation that the DMN, when analysed at the finer time scales, can be decoupled into two separate components. The anterior higher-order cognitive state includes the temporal poles (often associated with semantic integration; Tsapkini et al., 2011) and the ventromedial prefrontal cortex (typically implicated in emotion regulation and decision making; Fellows and Farah, 2007), exhibiting a strong connectivity with the posterior cingulate cortex (PCC), which is a key region of the DMN (Fransson and Marrelec, 2008). The posterior higher-order cognitive network encompasses the PCC/precuneus, bilateral superior and inferior parietal lobules, bilateral intraparietal sulci, bilateral angular and supramarginal gyri, and bilateral temporal cortex. These regions are classically associated with integration of sensory information, perceptual-motor coordination and visual attention, as well as processing of sounds, biological motion and theory of mind (Culham and Kanwisher, 2001).

**Fig. 2.**
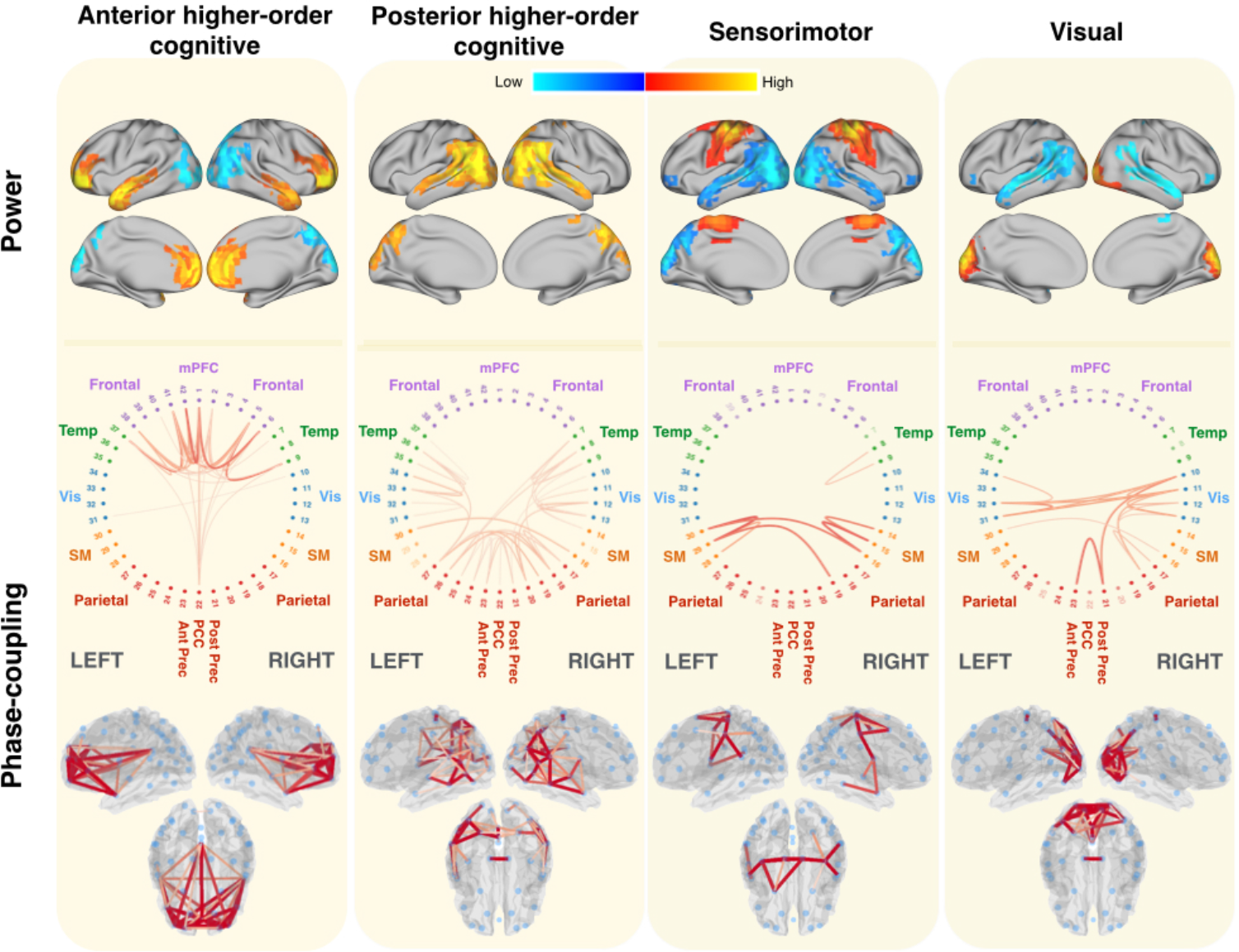
Brain states identified using Hidden Markov Modelling represent strong coherent (phase-coupling) networks, with strong increases in power tending to accompany increases in phase-coupling. Wideband (1-30Hz) thresholded power maps and phase-coupling are displayed for the two higher-order cognitive (anterior and posterior) states, and the visual and motor states. The two higher-order cognitive networks contain regions that suggest a subdivision of the default mode network. Power maps are relative to the temporal average, i.e. they are globally centred such that blue colours reflect power that is lower than the average over states, and red/yellow colours reflect power that is higher than the average over states. The coherence networks only show high-valued connections (see Methods for thresholding). In the circular phase-coupling plots, each numbered dot represents one brain region. **Fig. SI-1** shows the same information for the other eight states.

### Higher-order cognitive states have distinct spectral characteristics

Previous work looking at the global (temporally averaged) estimates of large-scale functional connectivity has demonstrated that different brain networks show correlation of power in different frequency bands (Hipp et al., 2012). Leveraging the fact that our model is spectrally resolved, we sought to investigate how power and phase-coupling varies with frequency in the different brain states.

For the four states shown in **Fig. 2**, **Fig. 3a** shows power versus coherence, with dots representing each brain region. These results are shown wideband (1-30Hz) and for three different frequency modes. The frequency modes were estimated following a data-driven approach (non-negative matrix factorisation, see Methods), which produced four frequency modes that approximately correspond to the classic delta/theta (0.5-10Hz), alpha (5-15Hz), beta (15-30Hz) and low gamma bands (30-45Hz). Possibly due to the relatively low signal-to-noise ratio in higher frequency bands, we found that the low gamma band mode did not exhibit any strong state-specific differences, and so we only show results for the delta/theta, alpha, and beta modes. Strong increases in power tended to (although not exclusively) accompany increases in coherence. Interestingly, the differences in coherence between the states are much more pronounced than the differences in power.

To expand on the specific spectral differences between the higher-order cognitive and the visual and motor states, **Fig. 3b** shows power and coherence averaged across all brain regions as a function of frequency. This shows frequency at full spectral resolution rather than the frequency modes used in the previous **Fig. 3a**. The anterior higher-order cognitive state is characterised by strong power and coherence in the slowest frequencies (delta/theta), whereas the posterior higher-order cognitive is dominated by the alpha frequency. More specifically, there is a strong component at ˜4Hz for the anterior higher-order cognitive state in both power and coherence, and a component at 10Hz for the posterior higher-order cognitive state also in both power and coherence. These results reveal that the two higher-order cognitive states, which may correspond to subdivisions of the DMN, exhibit more power and coherence than the visual and motor states. Moreover, they have very different dominant frequencies.

Given the key role attributed to the PCC and the medial prefrontal cortex (mPFC) within the DMN and resting-state networks more broadly (Fransson and Marrelec, 2008; van den Heuvel and Sporns, 2013), we next examined the state-specific frequency profile of the PCC and the mPFC to see if their spectral characteristics in the higher-order cognitive states are significantly different to the other (less cognitive) states. **Fig. 3c** shows the power in the PCC and the mPFC as well as the coherence between these two regions. The four considered states were compared to the global average (solid black lines; shaded areas represent the standard deviation across states), which corresponds to the power and coherence computed from a static (rather than dynamic) perspective. The PCC has more power across all frequencies in the posterior higher-order cognitive state, although the power in the slow frequencies for the anterior higher-order cognitive state is also significantly above the global average. By contrast, the mPFC shows high power in the anterior higher-order cognitive state, particularly in the delta/theta frequency range. Finally, global phase-coupling is high in the anterior higher-order cognitive state in delta/theta, whereas the posterior higher-order cognitive state exhibits high PCC connectivity in alpha and beta. Altogether, these results suggest that (i) the PCC has spectral properties that are unique to the higher-order cognitive states (consistent with the idea of the PCC being a hub region), (ii) the anterior higher-order cognitive state involves the PCC in slower frequencies than the posterior higher-order cognitive state for both power and phase-coupling connectivity, and (iii) these properties are only observed when we compute power and coherence specifically within the fast transient events that correspond to the HMM brain states, whereas the global, temporally averaged properties of the PCC (black line) are far less striking.

**Fig. 3.**
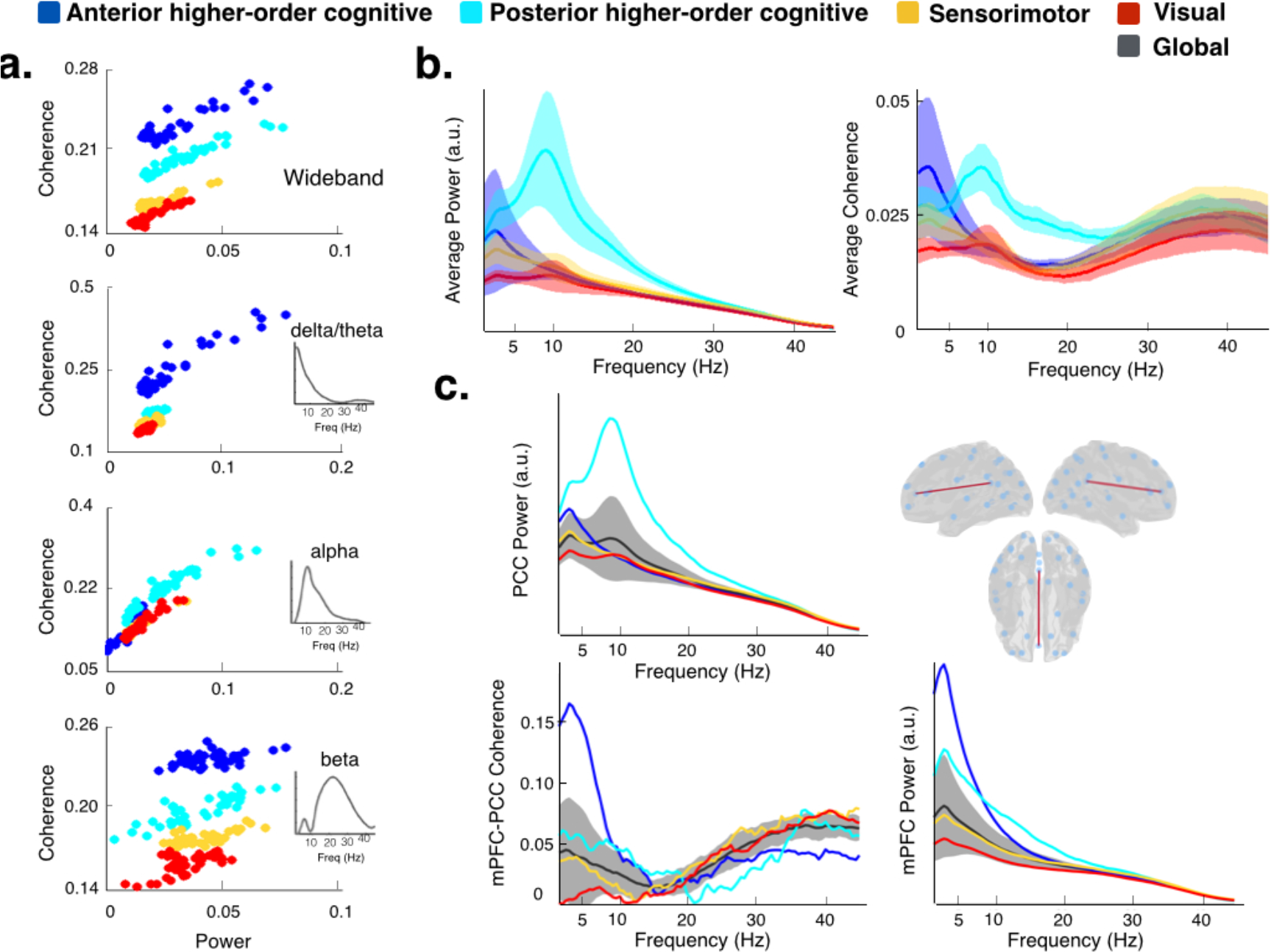
The two higher-order cognitive states have distinct spectral characteristics compared with the visual and motor states, including strong differences in PCC and mPFC activity. **(a)** Total connectivity of each region (defined as the sum of the values of coherence of the region with the rest of the regions) against power, for wideband and the three estimated frequency modes (see Methods), where each dot represent a different brain region. Both power and connectivity are higher for the higher-order cognitive than for the visual and motor states, with coherence exhibiting the largest difference. **(b)** Spectral profiles of the two higher-order cognitive (anterior and posterior) and the visual and motor states, in terms of power averaged across brain regions (left) and coherence averaged across all pairs of brain regions (right); shaded areas represent the standard deviation across brain regions (or pairs of regions). Fig. SI-4 shows the power spectra for the anterior/posterior precuneus alongside the PCC’s. **(c)** Power for PCC (top left), power for mPFC (bottom right) and coherence between mPFC and PCC (bottom left), for the four considered states in comparison to the grand average (black line, with the shaded areas representing standard deviation across states). The temporally average global power and coherence has a relative lack of spectral detail compared with the individual brain states.

We next examined the spatial distribution of these spectral differences using the frequency modes identified in Fig. 3. **Fig. 4** shows power and phase-coupling in brain space for each cognitive state and frequency mode (**Fig. SI-3** presents a similar view for the visual and motor states). This view clearly reflects that the two higher-order cognitive states have a distinct spatial distribution of power and connectivity. For example, we observe strong phase-coupling between frontal areas, mPFC and the PCC specifically in the delta/theta mode for the anterior higher-order cognitive state. By contrast, the posterior higher-order cognitive state is characterised by pronounced intraparietal and PCC/precuneus connectivity (specifically in the alpha frequency mode), and by some slow frequency power and phase-coupling in the temporal regions. As observed in **Fig. 2**, both power and connectivity exhibit strong interhemispheric symmetry.

**Fig. 4.**
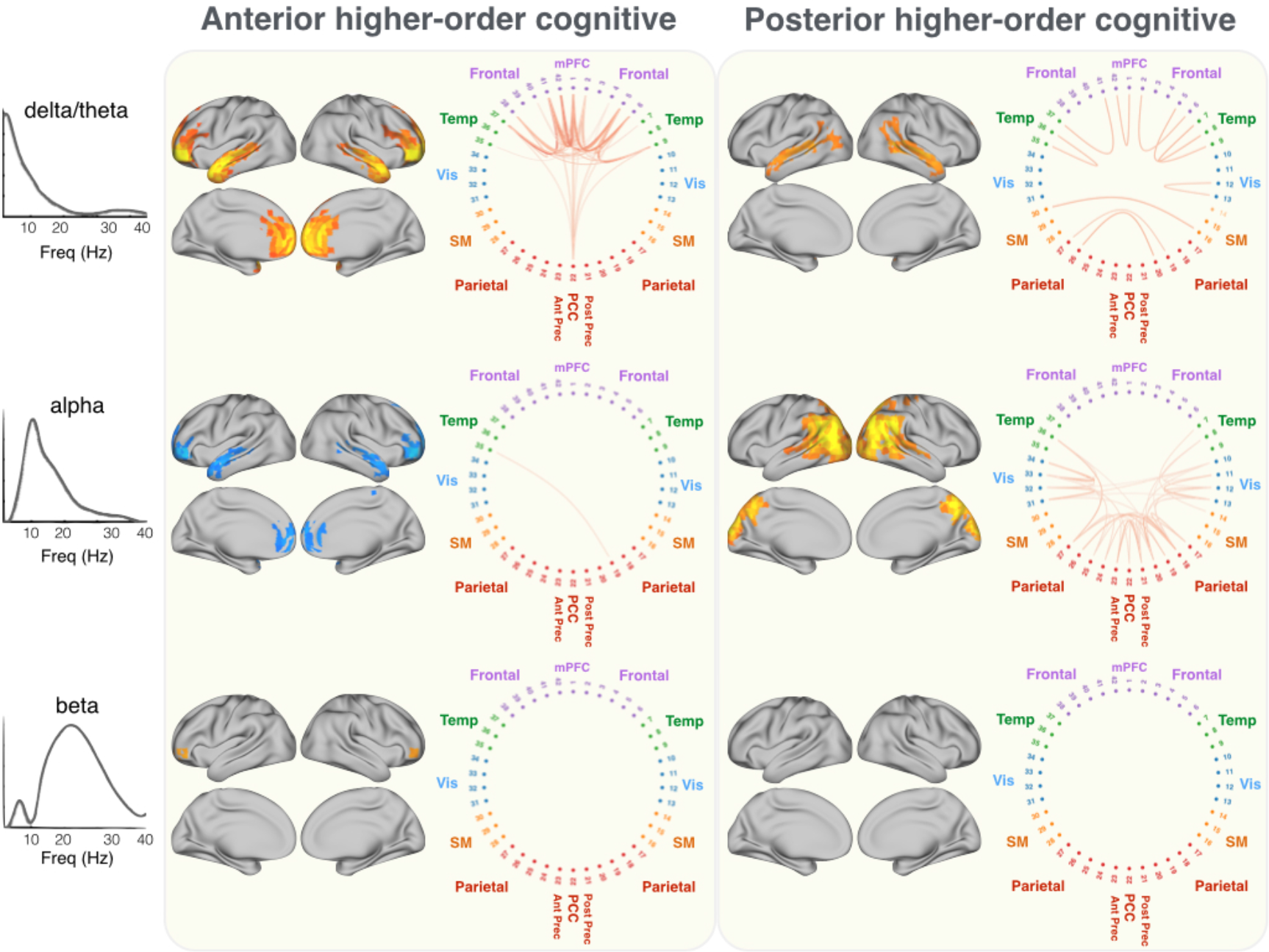
Frequency-specific relative power maps and phase-coupling (see Fig. 2 for details) for the anterior and posterior higher-order cognitive states, and for the three data-driven estimated frequency modes. Whereas activity (power and phase-locking) is dominant in the delta/theta frequencies for the anterior-cognitive state, the posterior-cognitive state is dominated by alpha. Both states exhibit strong phase-coupling with the PCC, but in different frequency bands. **Fig. SI-3** shows a similar view of the visual and motor states.

### Higher-order cognitive states have distinct temporal characteristics

Together with the state distributions, the HMM inference also estimates the time-courses of the visits to each of the brain states. We used these to look at the extent to which the temporal characteristics of the higher-order cognitive states differed to the visual and motor states.

**Fig. 5a** shows for each state: the dwell times (or life-times, i.e. the amount of time spent in a state before moving into a new state), the interval times between consecutive visits to a state, and the fractional occupancies (reflecting the proportion of time spent in each state). All HMM states were on average short-lived, their dwell times lasting on average between 50-100ms. (Note that, as shown empirically in the SI, phase-coupling can still be reliably measured for short state visits even at the slowest frequencies.) We observed longer dwell times for the posterior higher-order cognitive state than for the states that were not higher-order cognitive states (permutation testing, p-value<0.001). However, the largest differences are found in the interval times. Both of the anterior and posterior higher-order cognitive states have visits that are much more temporally separated than the other states (permutation testing: p-value<0.001 for both tests). The interval time distributions of the posterior higher-order cognitive state and, to a lesser extent, of the anterior higher-order cognitive state, have pronounced tails for higher interval times; as indicated by the mean of the distribution being much larger than the median. To further illustrate this, **Fig. 5b** shows the cumulative density function (CDF) of the interval times, which evaluates the proportion of intervals (Y-axis) that are longer than any given interval duration (X-axis). The CDF is particularly useful to examine the differences between the tails of the distributions. We observe that both higher-order cognitive states have significantly larger CDF values than the other states (significance for a confidence level of 0.01 is indicated by the lines on top of the panel, using permutation testing). For example, the time between state visits is longer than 1s, in ˜40% of the higher-order cognitive state visits, as compared to only ˜20% of the time for the other states. Importantly, this is not due to differences in fractional occupancy (depicted in the bottom panel of **Fig. 5a**), given that the fractional occupancies of the higher-order cognitive states are not significantly different from the visual and motor states. In summary, these results indicate that the higher-order cognitive states tend to last longer, but are not revisited for longer periods, than the visual and motor states.

**Fig. 5.**
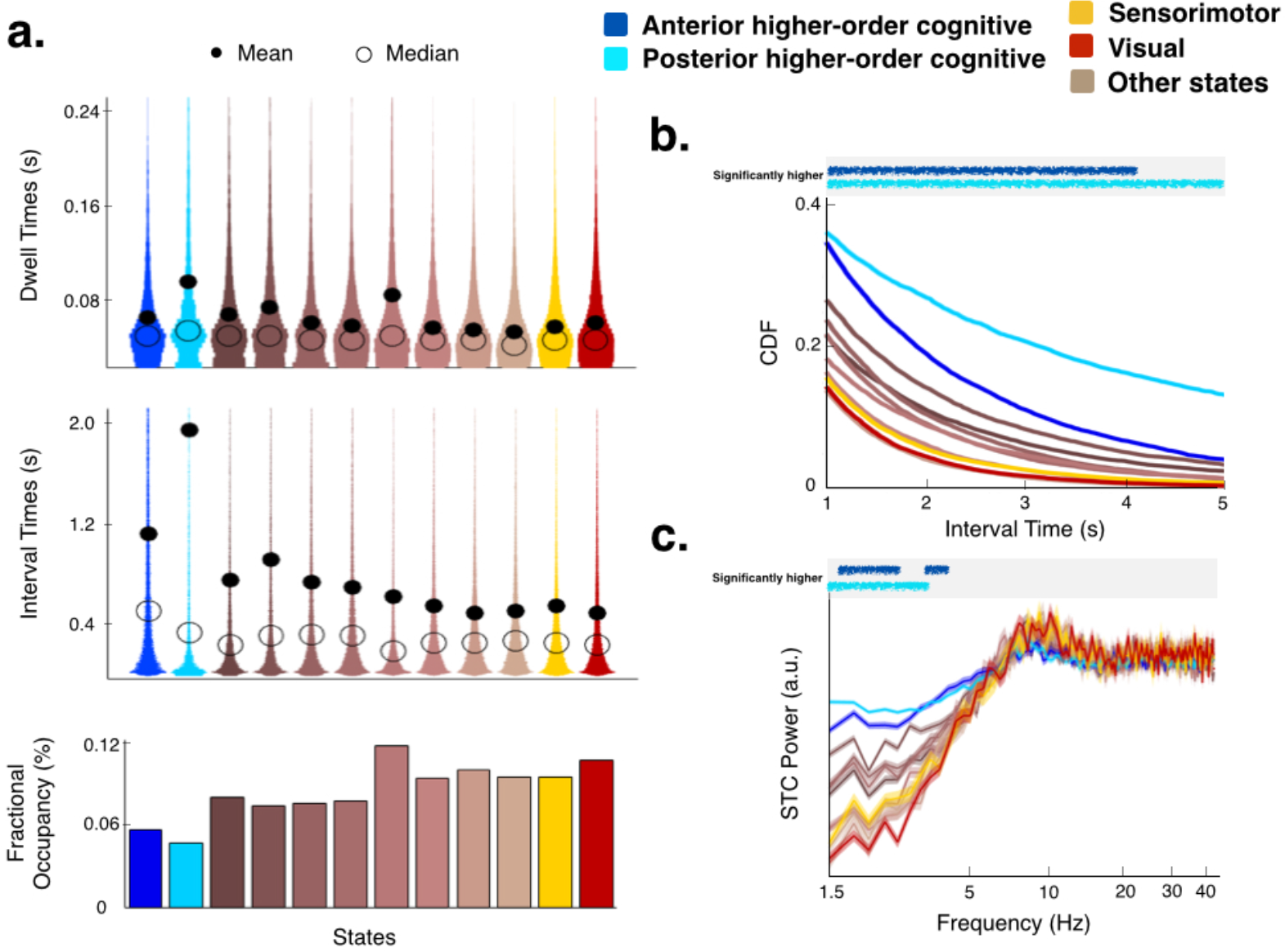
The higher-order cognitive states have distinct temporal features compared with the other states. The states depicted in brown colours are depicted in Fig SI-2, in an order that consistent with this figure (i.e. the first four are states with positive activation and the second four have a negative activation). **(a)** Distribution of state dwell times (time spent in each state visit), distribution of interval times between state visits, and fractional occupancies (proportion of time spent in each state). The dwell times are significantly longer for the posterior higher-order cognitive state than for all the other non-cognitive states (p-value<0.001), and the interval times are significantly longer for the two higher-order cognitive states than for the other states (p-value<0.001). **(b)** Cumulative density function (CDF) for the interval times, reflecting much larger tails for the interval time distribution of the two higher-order cognitive states; both of which have significantly larger CDF values than the other states (permutation testing; statistical significance for a confidence level of 0.01 is indicated by the lines on top of the panel). **(c)** Spectral analysis of a point process representing the onset of the state events, computed separately for each state (99% confidence intervals are indicated by shaded areas); no slow oscillatory modes in the state occurrences themselves is revealed. The higher-order cognitive states have a stronger power in the 1.5 to 5Hz range of frequency than the rest of the states (statistical significance using permutation testing is indicated on top, using a confidence level of 0.01).

We also investigated whether brain states are visited rhythmically, i.e. aligned to the peaks and troughs of a possible oscillation. Modelling the state events as a point process (where a state occurrence is defined at the onset of the state visit), we computed the spectra of these processes to find if there were any strongly characteristic frequencies. **Fig 5c** shows the power of the event point process for all states (shaded areas reflect 99% confidence intervals), which does not show any strong frequency mode and has particularly low power in the slow frequencies. This is in line with recent findings in task (Sherman et al, 2016), where it is only through trial-averaging (given the temporal variability of the task-related events within each trial), that these events appear as sustained oscillations. Also, consistently with the above results, we observed a higher power in between 1 and 5Hz for the higher-order cognitive states (significance for a confidence level of 0.01, using permutation testing, is indicated on top), reflecting the longer dwell and interval times for these states.

## Methods

### Data and preprocessing

As part of the UK MEG Partnership, 77 healthy participants were recruited at the University of Nottingham. A final cohort of 55 participants (mean age 26.5y, maximum age 48y, minimum age 18y, 35 males) was selected for analysis, discarding 22 subjects because of excessive head motion or artifacts. To avoid effects of tissue magnetisation, MEG data were acquired prior to participants entering the MRI. Resting-state MEG data were acquired using a 275-channel CTF MEG system (MISL, Coquitlam, Canada) operating in third-order synthetic gradiometry configuration, at a sample frequency of 1200Hz. MRI data, used here for the purpose of MEG coregistration, were acquired using a Phillips Achieva 7T system. (See Hunt et al., 2016 for further details about MEG and MRI acquisition). MEG data were then downsampled to 250Hz, filtered between 1 and 45Hz and source-reconstructed using LCMV beamforming (Van Veen et al., 1997; Woolrich et al., 2011) to 42 dipoles covering the entire cortex excluding subcortical areas (MNI coordinates are shown in **Table SI-1**). Bad segments were removed manually and correction for spatial leakage was applied using the technique described by Colclough et al. (2015). The effect of using alternative methods for leakage reduction is discussed in the SI.

### The Hidden Markov Model

As a general framework, the Hidden Markov model (HMM) assumes that a time series can be described using a hidden sequence of a finite number of states, such that, at each time point, only one state is active. In practice, because the HMM is a probabilistic model, the inference process acknowledges uncertainty and assigns a probability of being active to each state at each time point. Effectively, this amounts to having a mixture of models (or states) explaining the data at each time point, where the mixture weights are the state probabilities. Importantly, the probability of a state being active at time point *t* is modelled to be dependent on which state was active at time point *t-1* (i.e., it is order-one Markovian). The model then assumes that the data observed in each state are drawn from a probabilistic observation model. The observation distribution is of the same family for all states, whereas the observation model parameters are different for each state. The different varieties of the HMM are thus given by which family of probabilistic observation distribution is chosen to model the states. This is useful because different observation distributions can be adequate for different data modalities (Baker et al, 2014; Vidaurre et al, 2016; Vidaurre et al 2017) while preserving a common framework. This can facilitate integration of results across modalities. The variety of the HMM introduced in this paper is presented in the next section along with some theoretical and practical discussion about its properties. Whichever choice of the HMM state distribution, the model can be applied to each subject independently or to the concatenated data of all subjects, such that a group estimation of the states may be obtained. In this paper, the states were defined at the group-level; however, the information of when a state becomes active (i.e. the state time course) is still specific to each subject. Inference on the model (i.e. the estimation of the parameters of the posterior distribution) is carried out using variational Bayes (VB), a method providing an analytical approximation at a reasonable cost by assuming certain factorisations in the posterior distribution; we refer to (Vidaurre et al, 2016) for further details about the inference scheme. Still, because of the high sampling rate of our MEG data (250Hz) and the relatively high number of subjects (55), standard VB becomes both time and memory consuming. On these grounds, we used stochastic inference to further alleviate computation time (Vidaurre et al, 2017b) such that an average run would take approximately 5h using a standard workstation with manageable memory usage. After the inference process, the Viterbi path is computed; this is defined as the most probable sequence of (hard assigned, i.e. non-probabilistic) states, and can be analytically computed, given the current estimation of the state observation models, using a modification of the standard HMM state time course inference (Rabiner 1989).

### The embedded Hidden Markov Model

Here, we apply a novel variety of the HMM to *raw* (instead of power envelope; Baker et al., 2014) time-courses. This allows us to detect changes not only in power but also in phase-locking. Although this was already the case with the HMM-MAR (Vidaurre et al., 2016), the MAR observation model works optimally with a limited number of regions and does not scale to whole-brain analysis. In this approach, our definition of observation distribution describes the neural activity over a certain time window using a Gaussian distribution with zero mean (i.e. using the covariance matrix) to model the entire window; this is equivalent to saying that our observation model corresponds to the data *autocovariance* across regions (sometimes referred to as lagged cross covariance) within such window. For example, if we use a time window of 60s, having time point *t* assigned to a certain state means that, for our current 250Hz sampling rate and 42-region parcellation, the activity of the 42 channels over a window of 15 time points centered at *t* gets described by such state’s (15 × 42 by 15 × 42) autocovariance matrix. This multivariate autocovariance matrix can effectively capture patterns of linear synchronisation in oscillatory activity for those time points when a particular state is active, i.e. our model can describe state-wise phase-locking. This is mathematically equivalent to using a standard HMM with a Gaussian observation model on an “embedding” transformation of the original data (see **Fig. SI-1** for an illustration of the entire pipeline). In our case, with 55 subjects and 5min of data at 250Hz per subject, this amounts to running the HMM on a large (4125000 by 630) volume of data. As a result of the computational advantages of stochastic inference (Vidaurre et al, 2017b), it is still possible to handle such large amounts of data. However, it requires estimating (630 × 629) / 2 = 198135 parameters within the multivariate autocovariance matrix per state, which, above and beyond computational considerations, can lead to severe overfitting problems. To avoid this issue, we ran the HMM on a PCA decomposition of the “embedded” space. This not only greatly reduces the complexity of the state distributions but also naturally focuses the slower frequencies in the data. This is a consequence of PCA aiming to explain the highest possible amount of variance in the time series, in combination with the 1/f nature of electrophysiological data (i.e. that most of the power, or variance, is concentrated in the slow frequencies). In particular, we use twice principal components as the number of channels (i.e. 84 principal components). Note that given that the time-series from the source-space parcellation are orthogonal after leakage correction (Colclough et al, 2015), the PCA step can only leverage autocorrelations and non-zero lag cross-channel correlations to achieve an optimal decomposition. Since the non-zero lag cross-channel correlations are very small in comparison with the within-channel autocorrelation of the data (as shown in Vidaurre et al, 2016), we chose a number of PCA components that is a multiple of the number of channels; otherwise, because of the very nature of PCA, the “extra” PCA components will be explaining variance from just a few channels. Precisely which channels is mostly arbitrary, given that all channels were standardised to have the same variance. For example, for our 42-regions parcellation, using 100 PCA components will result in (100 − 42×2 =) 16 PCA components explaining variance from a (mostly) random subset of 16 regions.

### Source-reconstructed dipole ambiguity

It is an acknowledged issue that source-localised EEG and MEG data have an arbitrary sign as a consequence of the ambiguity of the source polarity. As source reconstruction, in this case through beamforming (Hillebrand and Barnes, 2005), is done for each subject separately, the sign of the reconstructed dipoles risks being inconsistent across subjects. This is not a problem when modelling the power time courses, but is a cause of concern for models based on the raw signal because connectivity between any pair of regions can cancel out at the group level if regions have their time courses flipped for a subset of the subjects. Here, we extend and generalise the basic idea in (Vidaurre et al, 2016) to multiple leakage-corrected channels, making the assumption that the lagged partial correlation between each pair of brain regions, across several different lags, has the same sign across subjects. (We choose to use partial correlation instead of simple correlation because this is a direct measure, i.e. there are no other channels interfering in the “sign relation” between every pair of channels). More explicitly, for all lags (for example, in between *α*=−10 and *α*=+10), we aim to find a combination of sign flips for each subject such that the function

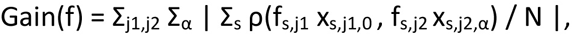

is maximised. Here, *s* cycles through the *N* subjects, *j1* and *j2* cycle through regions, *x*_*s,j1,α*_ represents the data time series for subject *s* and source *j1* that have been lagged *α* time points, *f*_*s,j1*_ takes the value −1 or +1 and represents whether channel *j1* is flipped for subject *s*, *ρ()* represents the partial correlation between a pair of time series, and ‖ is absolute value. The idea is that, provided the aforementioned assumption, *Gain(f)* will be maximised when the signs are correctly aligned. For example, if there is a strong genuine anti-phase relationship (leading to negative correlation) between a given pair of regions, the sign for these regions will be pertinently flipped for those subjects having an in-phase relation (leading to positive correlation) such that the negative correlations do not get partially cancelled out by the occasional positive correlations when averaging across subjects.

To find the best combination of sign flips such that *Gain(f)* is maximised is an integer programming problem, and, thus, finding an exact solution is NP-hard. The computationally expensive step is to compute the (no. of channels by no. of channels) partial correlation matrix for each subject and lag (in our data, 21lags × 55subjects matrix inversions of size 42regions by 42regions). Once this is computed, it is relatively inexpensive to evaluate the function *Gain(f)* given the equivalence

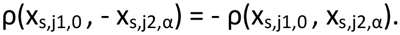

Therefore, we can afford to evaluate many different solutions, for example by multiple instantiations of a greedy algorithm with random initialisations of the signs. Although in this paper we limited ourselves to this simple approach, other more sophisticated search procedures can easily operate on this scheme.

### Extracting spectral information

Once the HMM has found the states on the basis of the data’s transient spectral properties, a logical further question is how can we extract and represent the states’ spectral properties in an informative way. We can extract the spectral information (power and phase-coupling) from the multivariate autocovariance matrix in each state’s observation model as it has a direct correspondence to the parameters of a MAR model (Lütkepohl, 2005), which contains the spectral information in the system. However, this estimation is biased towards the low frequencies due to the PCA dimensionality reduction step discussed in the previous section. So that we can effectively access high frequency information, we instead made use of the state-wise multitaper approach introduced in (Vidaurre at al, 2016), which will provide us with power and phase-locking coherence for each frequency bin (chosen to be in between 1 and 45Hz) and state without any PCA-induced bias.

Once we have estimated power and coherence for each state, we factorise this information into different components or frequency modes for ease of interpretation and visualization. We could use the traditional frequency bands for this matter, but instead we opted for estimating these in a data-driven fashion. To do this, we constructed a matrix by concatenating the spectrally defined coherence (the spectral feature that is more interesting for our purposes) across all states and pairs of regions. We shall denote this matrix as *A*. More specifically, *A* has (12 states × 861 pairs of regions =) 10332 rows and 90 columns, 90 being the number of frequency bins that we obtained from the multitaper analysis. We then applied a non-negative matrix factorisation (NNMF) algorithm (Berry et al., 2007) on *A*, asking for four components. NNMF aims to find a factorisation *A* = *WH*, where *W* has dimension (10332 by 4) and *H* has dimension (4 by 90), such that all the elements in *W* and *H* are positive. Each row of *H*, then, represents the spectral profile of this component, inferred from the data. These components turn out to roughly correspond to the canonical delta/theta, alpha, beta and (lower) gamma bands. Of these, our interpretations are focused on the first three components (displayed in **Fig. 4**, left panels), excluding the fourth (gamma band) component for being potentially less relevant to understanding large-scale synchronisation. Having four frequency modes allowed us, however, to have beta separated from gamma, providing a cleaner view on the data. With the component spectral profiles *H* in hand (referred to as frequency modes throughout the paper), it is straightforward to obtain values of coherence for each state, pair of regions and NNMF component. We do so by simply multiplying the respective (1 by 90) vector of coherence values by the corresponding transposed row of *H* (90 by 1). For power, we follow the same procedure, reusing the component spectral profiles that we computed for coherence. Wideband results (**Fig. 1**) correspond to a simple average across all frequency bins.

For purposes of visualization, in **Fig 1**, **Fig. 4** and **Fig. SI-1**, we showed only the functional connections that were the strongest in absolute value. To avoid setting an arbitrary threshold, we separately fitted, for each state and NNMF frequency mode (and wideband), a mixture of two Gaussian distributions to the population of functional connections, such that we only show the connections that have more probability of belonging to the Gaussian distribution representing the strongest connections. When the population of functional connections is well represented by a single Gaussian distribution, that is indicative that there are no connections that are pronouncedly stronger than the average connectivity within the state, in which case we do not show any.

## Discussion

We show that large-scale networks in resting-state magnetoencephalography can be well-described by repeated visits to short-lived transient brain states. Here, a state is defined as a distinct spatially-and spectrally-defined pattern of network activity across the set of considered regions, which span the whole brain. These patterns of activity and phase-coupling were found to be largely symmetric across hemispheres, and corresponded to plausible functional systems including sensory, motor and higher-order cognitive networks. Two higher-order cognitive brain states (or networks) contained regions suggesting a subdivision of the default mode network (DMN). These subdivisions operated in distinct frequency bands, with one state corresponding to a posterior network with high power and coherence in the alpha range (8-12Hz), and the other to an anterior network with high power and coherence in the delta/theta range (1-7Hz).

### Fast Transient Brain States and Slow Rhythms

In previous work on resting-state MEG, an HMM was used to identify fast transient brain states characterised by co-modulations in power (Baker et al., 2014). This approach could identify brain states that corresponded well with canonical resting-state networks in fMRI, and showed states switching on ˜100-200ms time scales. However, being based on band-limited power time courses, it was unable to identify potentially faster phenomena that are only apparent in the raw electrophysiological time-courses. By contrast, the approach presented in this paper can find brain states with distinct, brain-wide networks of spectrally resolved power and phase-locking from *raw* MEG time-courses. As a consequence, states were found to switch on ˜50-100ms time scales, revealing fast dynamic power and phase-locking information not apparent from a static perspective. Being able to identify phase-coupling is crucial, as this has been proposed as an important mechanism for regulating the integration and flow of cognitive content (Fries, 2005; Fries 2015; Engel et al., 2013). The identification of large-scale networks of phase-locking in the present work is consistent with the idea that the brain spontaneously evokes the same network dynamics that we see in task (Smith et al., 2009).

But, how can the fact that state visits are often under 100ms in duration be compatible with the slow frequencies (e.g. delta/theta bands) that characterise the states? For example, an 8Hz theta cycle, which is in the realms of the frequencies reported here, has a period of 125ms. This can be reconciled by noting that, although we do not in general capture prolonged oscillations, spectral estimation does *not* actually require entire cycles. Unlike sliding window approaches, the HMM provides a large number of separated sub-cycle wave segments, with which the spectral estimation at the slow frequencies is possible. This is because frequency is defined instantaneously, and depends on the gradient on the signal (Huang et al., 2009), which is theoretically defined at each time point. We have performed simulations to prove this empirically (see **SI** and **Fig. SI-6**).

### Subdivision of the Default Mode Network

Our results identified two higher-order cognitive states or networks that showed particularly high power and coherence in comparison with the other states. These higher-order cognitive networks also exhibited different temporal dynamics in their state occurrences, notably with longer periods of times between state visits. One of these states represents a posterior network including PCC, precuneus and bilateral intraparietal regions. The other encompasses anterior areas including mPFC and temporal poles, exhibiting strong connectivity with the PCC. Consistently with previous studies in fMRI (Smith et al., 2012), these results afford the interpretation of the DMN being separable into anterior and posterior subdivisions. The present work, however, offers an important new insight into the electrophysiological properties of these distinct subnetworks. Here, the two subdivisions are distinguished from each other by operating within very different frequency bands: in the alpha band for the posterior network, and in the delta/theta band in the anterior network. Furthermore, the PCC may be acting as a link between the two higher-order systems, given that it is present in both networks and has particularly strong delta/theta band connectivity with the mPFC in the anterior higher-order cognitive state. This is consistent with previous work where, based on band-limited power correlations yet ignoring phase-coupling, the PCC was proposed to serve as a hub (de Pasquale et al., 2012).

The operation of these large-scale cortical phase-coupling networks in very different frequency bands may reflect the different intrinsic timescales that they specialise in within the temporal domain. Spike count autocorrelograms from single neurons vary across region, with parietal/prefrontal regions showing exhibiting shorter/longer timescales respectively (Murray et al., 2014). This can be accounted for using a model that accounts for variation in both long-range connectivity and local circuit dynamics (Chaudhuri et al., 2015), and such properties may explain variation in ‘temporal receptive windows’ across brain regions within temporally extended tasks (Lerner et al., 2011). How these within-region properties may interact with changes in interregional phase-coupling during tasks remains unclear, and is an important area for future investigation.

### Posterior Cingulate Cortex in Resting-state MEG

Despite having been attributed a key role in the resting-state, and in particular in the DMN (Buckner et al., 2008; Fransson and Marrelec, 2008), the PCC has been somewhat under-represented in the resting M/EEG literature, possibly due to the relatively low signal-to-noise ratio (and, hence, visibility) in M/EEG (Hillebrand and Barnes, 2002). For example, previous analyses of resting MEG data reported putative DMN networks that did not include the PCC (Brookes et al., 2011, Baker et al., 2014; Hipp et al., 2012). One possible reason is that the PCC’s role as a hub potentially involves many different network states, in such a way that is supressed when examining *differences* between networks or states (Baker et al., 2014). Notably, in a resting MEG study that used time-windows of high band-limited power correlation between nodes of the DMN, the PCC exhibited the highest of these correlations (de Pasquale et al., 2012). Here, the PCC is highly visible when networks are characterised by phase-coupling, especially in the anterior and posterior higher-order cognitive states.

Given the issues in representing the PCC in MEG, it is often merged with precuneus. However, these are remarkably different regions, with different structural connectivity profiles and distinct functional roles (Leech et al., 2011). Here, we used a parcellation that separated the PCC, the anterior precuneus and the posterior precuneus. **Fig. SI-4** shows power and phase-coupling with mPFC for each state and each of the three regions. Some differences can be clearly recognised between regions and, in particular, between the PCC and the two precuneus regions. Remarkably, coherence with mPFC is three times higher for the PCC than the precuneus (both anterior and posterior) in the anterior higher-order cognitive state. Also, in the anterior higher-order cognitive state the PCC exhibits strong activity in the delta/theta frequency mode whereas the precuneus does not.

### Relationship between Power and Coherence

The approach we use identifies brain states characterized by distinct power and phase-coupling. We find that states that show increase in power, often, although not exclusively, also show increases in coherence. However, it is well known that, due to changes in the signal-to-noise ratio, increases in power can augment the estimated coherence even in the absence of an actual change in the interactions between the regions. The same phenomena can cause increases in variance to effect correlation-based measures of functional connectivity (Cole et al., 2016; Duff et al., 2017). Therefore, some of the observed changes in coherence between states might be caused by differences in power. Notably, while we find that power and coherence are generally positively correlated, there are various aspects of phase-coupling that cannot be explained by changes in power. For example, the differences between the states are much stronger in coherence than in power (see, e.g., **Fig. 3a**). Also, whereas the PCC has slightly more (low-frequency) power in the anterior higher-order cognitive state than in the other states, phase-coupling is the feature most strongly stands out from the rest (see **Fig. 3c**).

### Gamma-band

This study focused on lower frequency bands (1-45Hz). Because of the methodological considerations discussed in the Methods section, and because of the higher signal-to-noise ratio in lower frequency bands (1-30Hz), low Gamma frequencies (30-45Hz) did not reveal any clear state-specific differences. However, we would expect there to be different patterns in gamma, given the possible top-down modulation of these frequencies by the slow frequencies (Canolty et al., 2006). Because of the crucial importance of gamma in cognition, having a key role in information transference between regions and plasticity (Martínez et al., 1999; Miltner et al., 1999; Mehta et al., 2002; Fries et al., 2007; Buzsáki and Draguhn, 2004; Fries 2015), it is of primary interest to understand how gamma frequency is modulated at the whole-brain level across different states. This will be an important area for future studies.

### Number of Brain States

A central parameter in our approach is the number of HMM states. Here, we have chosen it to be twelve. Of note, we do not claim this number to be closer to any biological ground-truth than, for example, eight or sixteen. Although it is possible to guide the choice of the number of states using quantitative measures like the free energy (Vidaurre et al., 2016), or even using non-parametric approaches that automatically determine the number of states (Beal et al., 2002), different numbers of states in practice just offer different levels of detail of brain dynamics. Indeed, examining different degrees of “abstraction” can itself reveal useful insights. For example, when we ran the proposed approach with six states (see **Fig. SI-5a**) the posterior higher-order cognitive state was fused with the SM/precuneus and VIS/precuneus states (depicted in **Fig. SI-2**). This is unsurprising considering their relatively similar spectral and spatial features. In this analysis, the left and right temporal states (see **Fig. SI-2**), which are characterised by high asymmetry and are possibly related to language, were also merged into a single symmetric state containing the patterns of both. In summary, running the HMM with different numbers of states and combining the results in a principled way can provide a hierarchical view of the data that is hidden to other approaches. Thanks to the stochastic scheme of inference (Vidaurre et al., 2017b), HMM runs are not computationally expensive to produce, facilitating these exploratory analyses.

### State exclusivity

The model specification of the HMM assumes that only one state is active at each point in time. However, the HMM inference actually assigns a probability to each state at each time point, so the strict exclusivity assumption is effectively relaxed in the estimation. Further, it is possible for network multiplexing to be expressed in the correlation of state fractional occupancies at slower time scales. Addressing the information contained in the state time courses at multiple time scales is an important area for future investigations.

### Summary

We have proposed an analysis approach that allows the investigation of dynamic changes in whole-brain phase-coupling in the resting state. Our study revealed that at these fast time scales, higher-order regions within the default mode network dissociate into two spatially, temporally and spectrally distinct states. These states potentially index different higher-order cognitive processes that themselves operate at different timescales. Although we have focused on this particular aspect of the data, the wealth of information contained in the model output opens many avenues for future analyses, hypotheses and questions. These include the dynamics of specific phase relations between areas, the whole-brain dynamics of gamma at rest, and the existence of changing patterns of communication between processes operating at different frequencies.

## Acknowledgements

This study was supported by the NIHR Oxford Health Biomedical Research Centre, an MRC UK MEG Partnership Grant (MR/K005464/1), a James S. McDonnell Foundation Understanding Human Cognition Collaborative Award (220020448), and the NIHR Oxford Health Biomedical Research Centre. The Wellcome Centre for Integrative Neuroimaging is supported by core funding from the Wellcome Trust (203139/Z/16/Z). DV is supported by a Wellcome Trust Strategic Award (098369/Z/12/ Z). MWW is supported by the Wellcome Trust (106183/Z/14/Z) and the MRC UK MEG Partnership Grant (MR/K005464/1). LTH is supported by a NARSAD Young Investigator Grant from the Brain and Behavior Foundation. ACN is supported by a Wellcome Trust Senior Investigator Award (ACN) 104571/Z/14/Z. BAEH is funded by an MRC Partnership Grant MR/K005464/1, and an MRC Doctoral Training Grant MR/K501086/1. Data were collected in Nottingham in the context of the MRC-funded MEG-UK partnership.

## Supplementary Information

### Slow-frequency spectral properties within fast state visits

We obtained state visits that were often well under 100ms (**Fig. 5**). How can this be compatible with the slow frequencies (e.g. delta/theta bands) that characterise the states? Here, we show that this is theoretically and practically possible through simulations. We have simulated data where segments of an 8Hz theta wave are interspersed with unstructured signal. We have performed three sets of simulations. In each of them, the duration of the wave segments (which are selected at random points of the theta period) are sampled from a Poisson distribution with mean 0.025s, 0.05s, and 0.1s, respectively. The separation between segments is sampled from a Poisson distribution with mean 1s in all cases, which makes the different wave occurrences to be completely phase-independent. Small-variance Gaussian noise is added to the generated signals. We simulated 20min of data at 250Hz for each simulation, and assumed a state time course that is active only at the time of the wave segments occurrences. Hence, the duration of the wave segments corresponds to the duration of the state visits. We then used the state-wise multitaper used in Results and in (Vidaurre et al., 2016) to assess the spectral content of the signal. **Fig. SI-6** shows the spectral estimation on the top, and an example of a wave segments for each mean dwell time in the bottom. In the three cases, and despite the short state visits, the state-wise multitaper is able to find the correct frequency, even when the frequency resolution is degraded somewhat as we make the wave segments (state visits) shorter, leaking power toward faster frequencies.

### Leakage reduction and phase-locking coherence estimation

In this work, we used the method proposed by Colclough et al. (2015) in order to reduce the effect of signal leakage (volume conduction). Without this step, the estimation of phase-locking gets dominated by a pattern of artefactual local connections that is common to all states. While the Colclough et al. approach has been shown to work well in the context of MEG amplitude correlations [Colclough et al. (2015)], its application for phase-locking networks is less well established. Recently, Pascual-Marqui et al. (2017) have challenged Colclough et al.’s approach, particularly within the context of estimating phase-locking measures, showing that under certain conditions, artefactual connections may arise. The authors also provide an alternative approach based on the multivariate autoregressive model that may overcome these issues.

To check the Colclough et al. approach in the context of phase-locking, we also applied the Pascual-Marqui’s approach on our real data, and compared the resulting state-specific phase-locking with the estimations depicted in **Fig. 2**. In **Fig SI-7**, phase-locking connectivity is shown for the same four HMM states, after applying Pascual-Marqui’s method. As observed, the differences between **Fig SI-7** and **Fig. 2** are limited, with the method proposed by Pascual-Marqui and colleagues being slightly more conservative. The main features of the HMM states are however preserved.

**Table SI.** List of regions used in the analysis, with their MNI coordinates. Numerical indexes correspond to those shown in Fig. 2, Fig. 4, and Fig. SI-2.

| Index | Area | MNI Coordinates (X,Y,Z) |
| --- | --- | --- |
| 1 | Medial PFC | (2,50,0) |
| 2 | Right Frontal Lobe | (42,34,16) |
| 3 | Right Frontal Lobe | (26,10,56) |
| 4 | Right Frontal Lobe | (18,42,40) |
| 5 | Right Frontal Lobe | (42,50,0) |
| 6 | Right Frontal Lobe | (26,50,24) |
| 7 | Right Temporal Lobe | (58,-22,8) |
| 8 | Right Temporal Lobe | (58,-46,0) |
| 9 | Right Temporal Lobe | (50,10,-24) |
| 10 | Right Visual Cortex | (10,-94,24) |
| 11 | Right Visual Cortex | (26,-94,8) |
| 12 | Right Visual Cortex | (50,-70,8) |
| 13 | Right Visual Cortex | (42,-78,-8) |
| 14 | Right Sensorimotor Cortex | (58,-6,32) |
| 15 | Right Sensorimotor Cortex | (42,-22,56) |
| 16 | Right Sensorimotor Cortex | (10,-30,72) |
| 17 | Right Parietal Lobe | (18,-70,56) |
| 18 | Right Parietal Lobe | (34,-78,40) |
| 19 | Right Parietal Lobe | (-54,-46,40) |
| 20 | Right Parietal Lobe | (50,-70,16) |
| 21 | Posterior Precuneus | (-6,-70,32) |
| 22 | Posterior Cingulate Cortex | (2,-46,24) |
| 23 | Anterior Precuneus | (2,-54,48) |
| 24 | Left Parietal Lobe | (-22,-62,56) |
| 25 | Left Parietal Lobe | (-38,-78,40) |
| 26 | Left Parietal Lobe | (58,-46,40) |
| 27 | Left Parietal Lobe | (-46,-70,16) |
| 28 | Left Sensorimotor Cortex | (-54,-6,32) |
| 29 | Left Sensorimotor Cortex | (-46,-22,56) |
| 30 | Left Sensorimotor Cortex | (-6,-30,72) |
| 31 | Left Visual Cortex | (-14,-94,24) |
| 32 | Left Visual Cortex | (-22,-94,8) |
| 33 | Left Visual Cortex | (-46,-70,8) |
| 34 | Left Visual Cortex | (-38,-86,0) |
| 35 | Left Temporal Lobe | (-62,-22,8) |
| 36 | Left Temporal Lobe | (-62,-46,0) |
| 37 | Left Temporal Lobe | (-46,10,-24) |
| 38 | Left Frontal Lobe | (-46,34,16) |
| 39 | Left Frontal Lobe | (-22,10,56) |
| 40 | Left Frontal Lobe | (-14,42,48) |
| 41 | Left Frontal Lobe | (-38,50,0) |
| 42 | Left Frontal Lobe | (-22,58,16) |

**Fig. SI-1.**
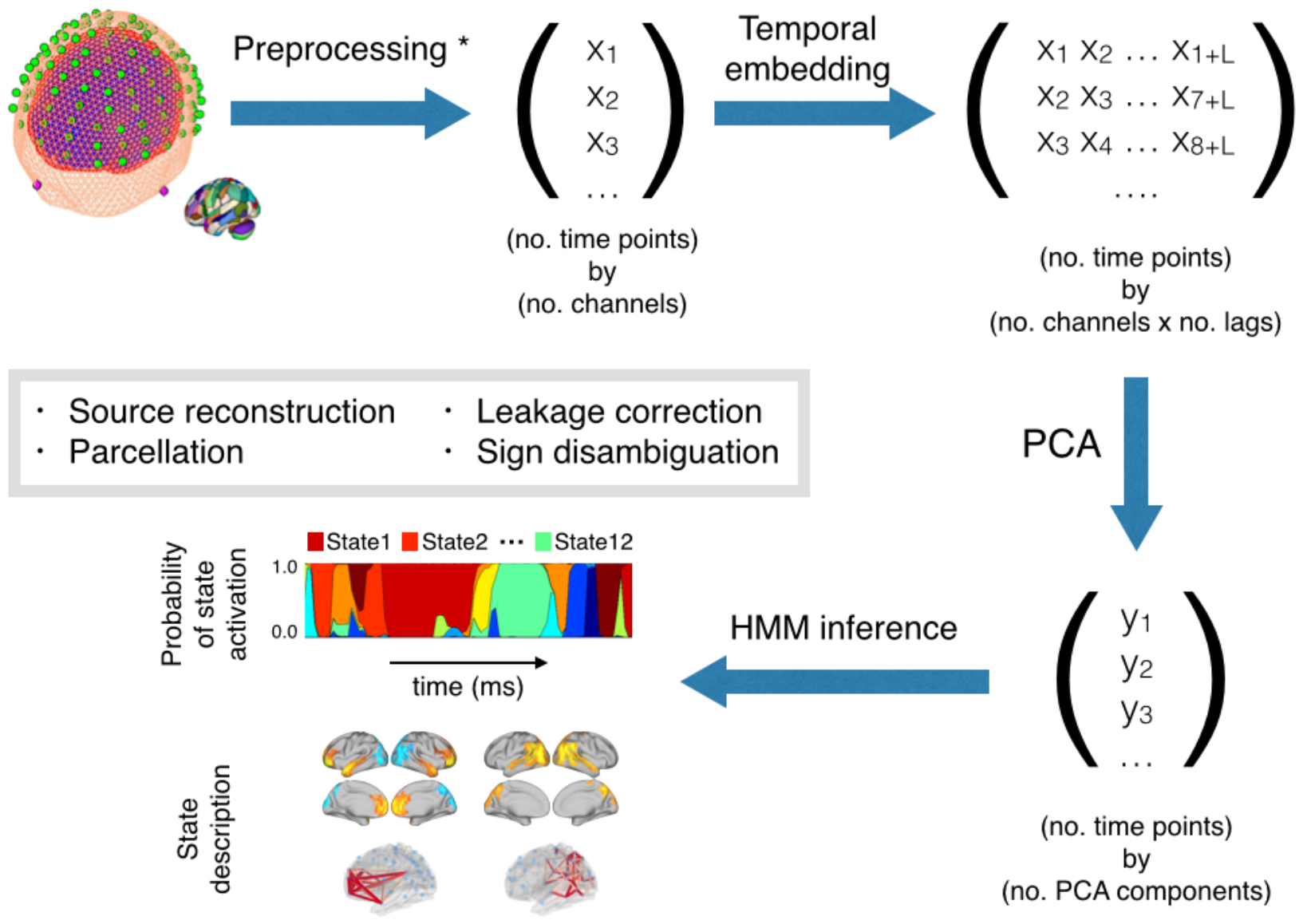
Schematic overview of the proposed method. (Related to Fig. 1). After preprocessing (source reconstruction, parcellation, leakage correction and sign disambiguation – see Methods), the data channels *X* are temporally-embedded (using *L* lags), PCA is applied for dimensionality reduction (producing PCA components *Y*), and HMM inference is then used to find the state time courses and the state parameters.

**Fig. SI-2.**
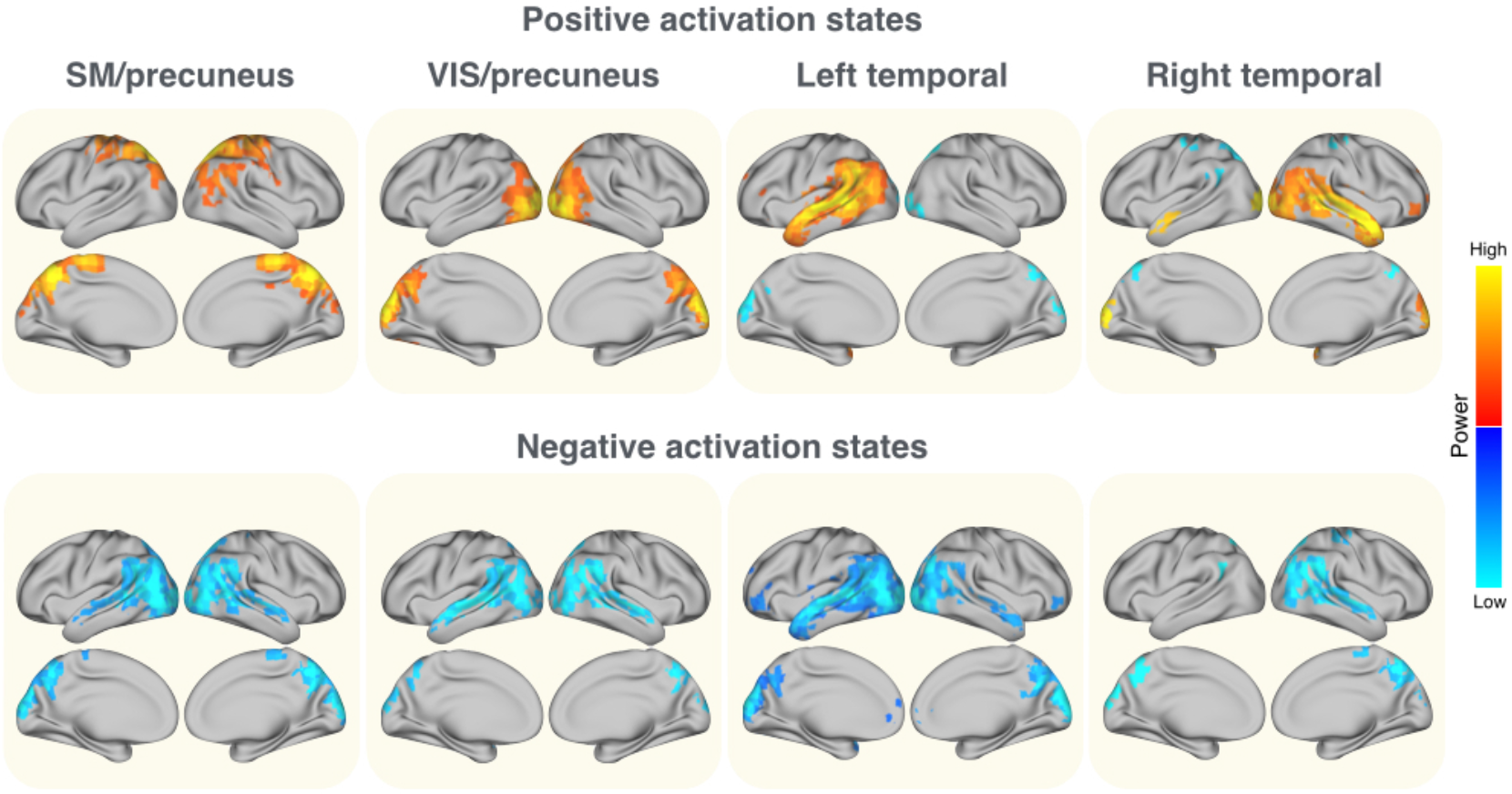
The remaining eight states of the HMM model, four of which (in the bottom) correspond to depressed (with regard to average) power and connectivity. (Related to Fig. 2)

**Fig. SI-3.**
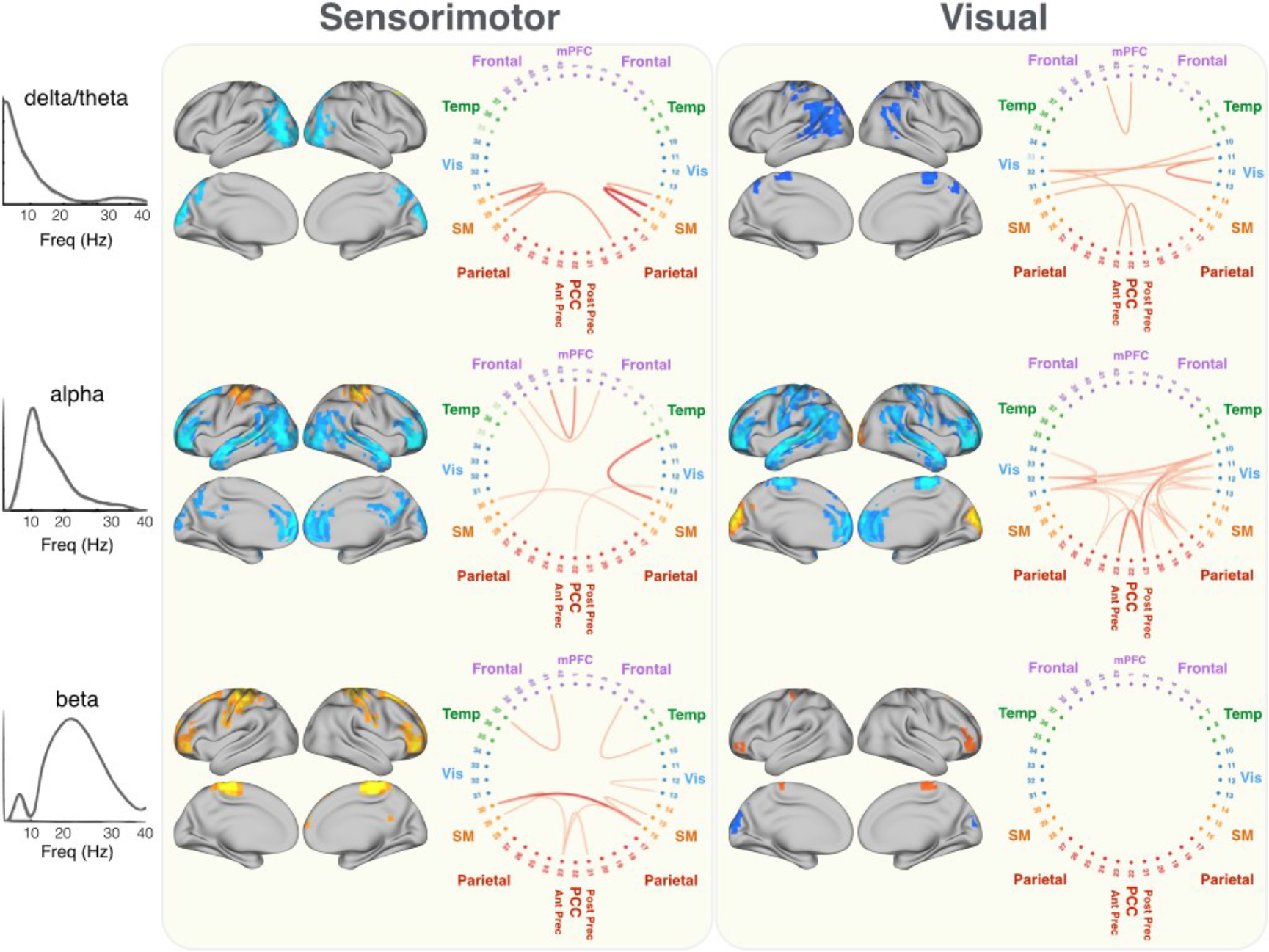
Frequency-specific power and phase-locking connectivity for the visual and motor states, for the three data-driven estimated frequency modes (see Methods). (Related to Fig. 4). With regard to connectivity, only the connections with highest absolute value are shown (see Methods).

**Fig. SI-4.**
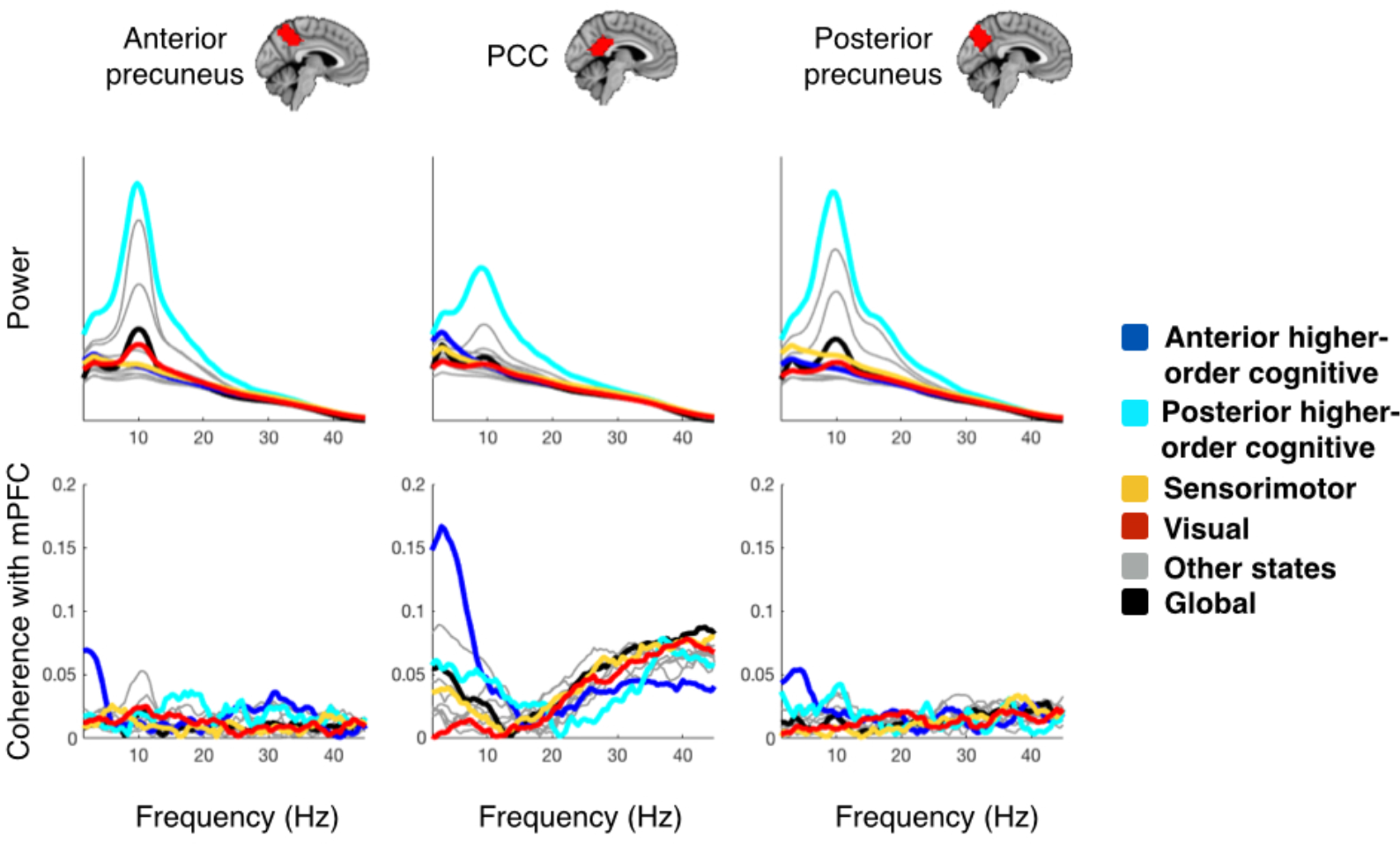
The spectral profile of the three regions is distinct when considered state by state. (Related to Fig. 3). The top three panels represent power as a function of frequency for the PCC and the two precuneus regions, separated by state and including the global power. The bottom panels reflect connectivity with the mPFC region.

**Fig. SI-5.**
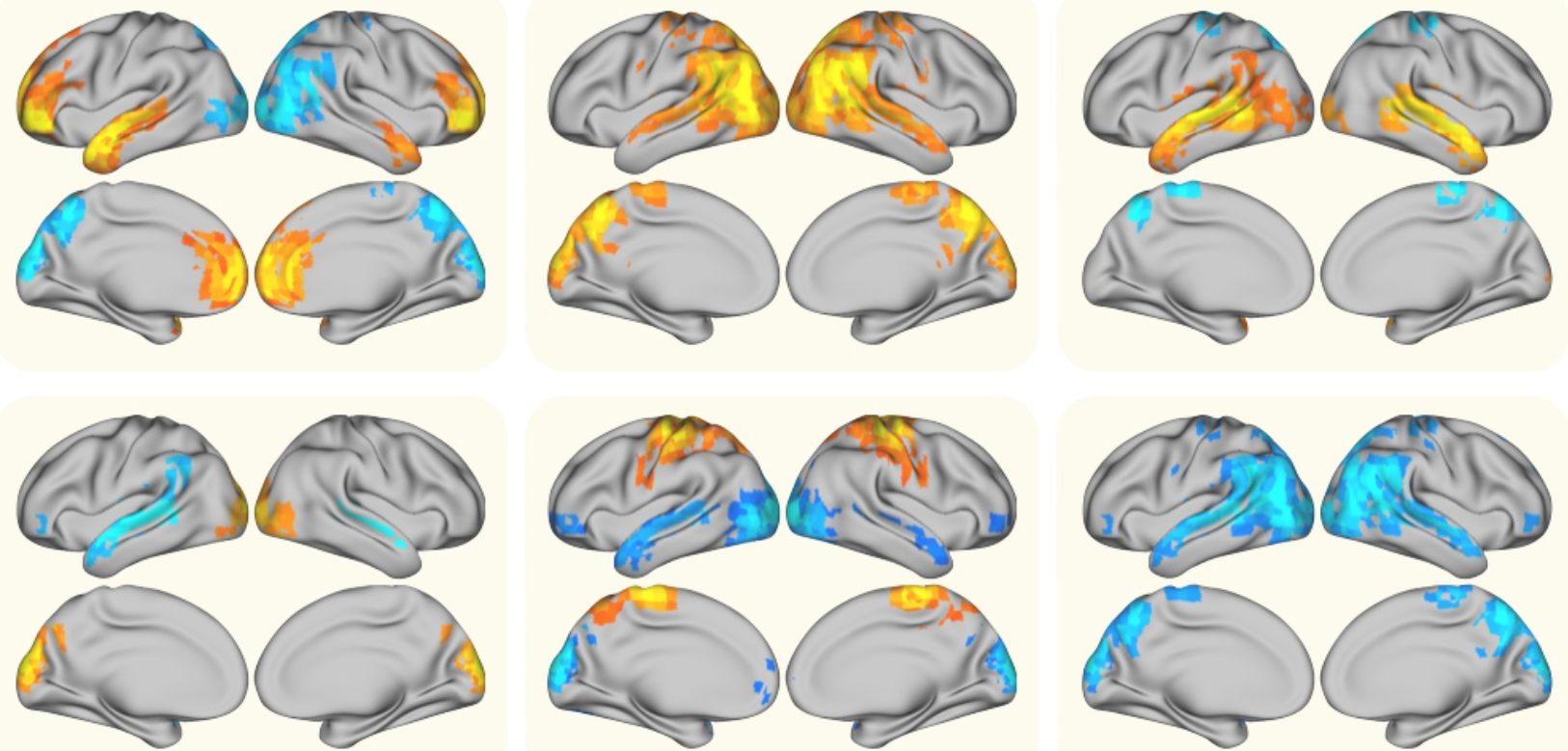
Spatial power maps for an HMM run with 6 states, where some states from the original 12 state analysis are now fused into fewer states.

**Fig. SI-6.**
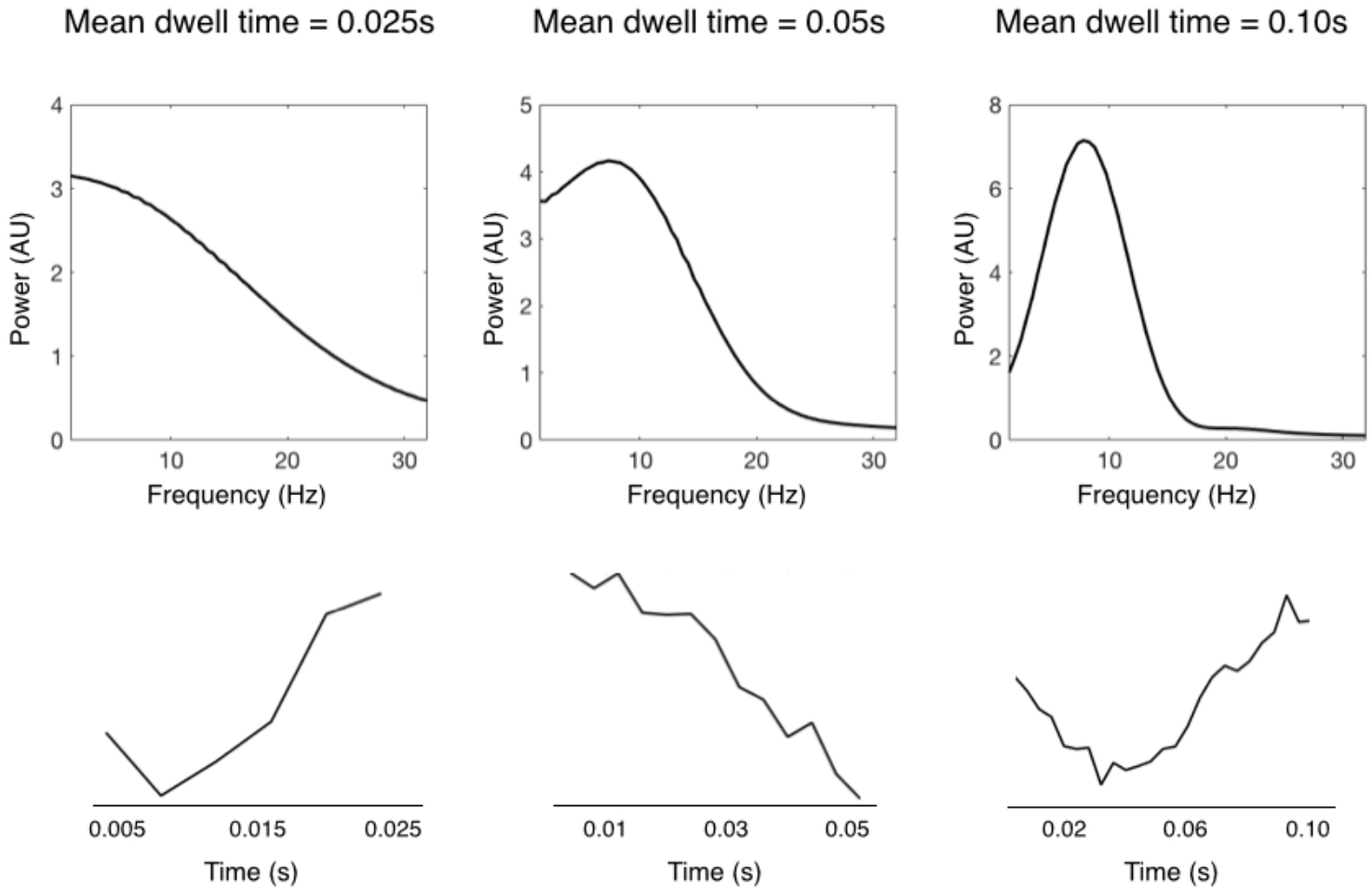
Spectral information from a collection of signal segments extracted from a canonical theta oscillation (8Hz), where the segments are shorter than the theta period (0.125s). The length of the segments has either mean 0.025s (left), or 0.05s (middle), or 0.10s (right); examples of the segments are shown in the bottom for each case. Despite the brevity of the segments, the spectral information of the underlying theta wave is correctly calculated.

**Fig. SI-7.**
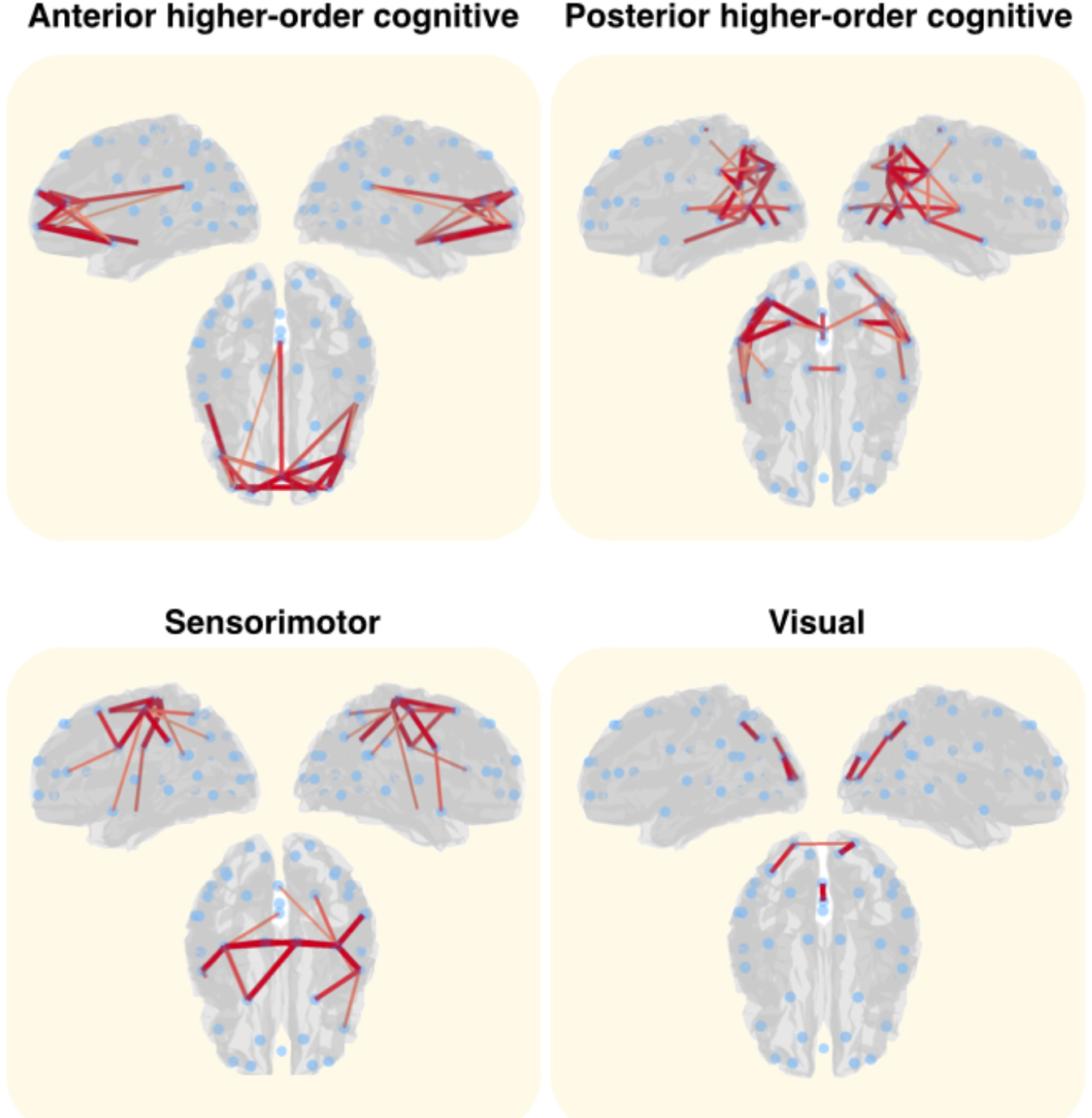
Phase-locking connectivity for the two higher-order cognitive (anterior and posterior) states, and the visual and motor states, when the method for leakage correction by Pascual-Marqui et al. (2017) is applied instead of the Colcough et al.’s method used in this work. The differences between the two methods are not large, with slightly fewer connections for Pascual-Marqui et al.’s method.

